# Dysbiosis of the larval gut microbiota of *Spodoptera frugiperda* Strains feeding on different host-plants

**DOI:** 10.1101/2022.05.25.493308

**Authors:** Nathalia Cavichiolli de Oliveira, Fernando Luis Cônsoli

## Abstract

The gut microbiota plays important roles in the bioecology of insects, including host plant adaptation and speciation. *Spodoptera frugiperda* has two well-established host-adapted strains with marked differences at the genetic and host plant utilization levels. We investigated whether differences in the gut microbiota would occur between the “corn” (*CS*) and “rice” (*RS*) strains of *S. frugiperda* when feeding on different crops. The gut microbiota of larvae fed on corn and millet was predominantly represented by *Firmicutes* followed by *Proteobacteria*, with an opposite pattern in larvae fed on cotton. No differences were observed between the *CS* and *RS* using PERMANOVA. PCoA analyses resulted in distinct bacterial clusters based on the host plant. Comparisons of strains gut microbiota at the phylum level resulted in differences only for larvae fed on cotton, but differences in the relative abundance of minor representatives at the genus level between strains were observed regardless of the food source used. We also found differences in the potential functional contribution of bacteria between the strains. In conclusion the gut microbiota of *S. frugiperda* is strongly modulated by the host plant while strains seemed to play a minor role in changing the abundance of members of the gut bacterial community.

## Introduction

It is increasingly recognized that microbes, particularly bacteria, play a crucial role in a wide range of aspects of the host physiology, ecology, and evolution (Bourtzis and Miller 2003; Gilbert et al. 2012). Thus, it is expected that hosts harboring beneficial microbes will have advantages over their peers, and their joint exposure to processes of natural selection will also act on characteristics that will contribute to select the most suitable microbiota to deliver the services required by the host, maintaining therefore the established association (Shapira 2016). The best symbiont-service providers will increase the fitness of their host and of their own, providing a basis for coevolution (Shapira 2016). Thus, the host-microbiota coevolution predicts the existence of species-specific gut microbiota composed of beneficial microbes adapted to the host (Shapira 2016).

The insect gut may harbor a diverse and abundant microbial community. The composition of the gut microbiota is prone to variations depending on the existence of specialized structures in the gut, gut pH, redox conditions, digestive enzymes, antimicrobial peptides, food type, and host habitat among others (Ryu et al. 2008; Yun et al. 2014). The gut microbiota can contribute with food digestion through the synthesis and release of digestive enzymes (Anand et al. 2010; Krishnan et al. 2014), and the detoxification of plant allelochemicals and synthetic insecticides (Kikuchi et al. 2012; Adams et. 2014; Almeida et al. 2018). Moreover, the gut microbiota can provide the host with vitamins and essential amino acids (Douglas 2006; Nikoh et al. 2012) as well as recycle waste nitrogen (French et al. 1976; Ohkuma et al. 1996) allowing the host to establish new associations with suboptimum food sources. In addition to their nutritional contributions, gut microbes can influence the process of species differentiation. Gut microbes of *Drosophila melanogaster* interfere with the mating choice as they influence the hydrocarbon composition of the cuticle that serve as contact sex pheromones (Sharon et al. 2011). Gut microbes were also shown to influence species differentiation by inducing hybrid lethality in the parasitic wasp *Nasonia* (Brucker and Bordenstein 2013). The fall armyworm *Spodoptera frugiperda* (Smith) (Lepidoptera: Noctuidae) is a severe, widespread, and well-known agricultural pest that was restricted to the Americas (Jonhson 1987) but has recently invaded the Old-World through Africa (Goergen et al. 2016; Otim et al. 2018) and has now reached the far east Asia (Liu et al. 2019; Padhee and Prasanna 2019). There are two well-characterized host-adapted strains of *S. frugiperda* identified so far that regardless of the genetic differences (Dumas et al 2019; Gouin et al. 2017) and postzygotic mechanisms of reproductive isolation (Kost et al. 2016) are still defined as strains carrying different bioecological traits belonging to a single species. Molecular dating analyses indicate these strains diverged more than 2 Myr ago (Kergoat et al. 2012). The two strains are identified as the rice (*RS*) and the corn (*CS*) strains, and they show a high level of genetic differentiation (Gouin et al. 2017), with differences in host plant utilization (Veesntra et al. 1995; Pashley et al. 1995). The *CS* feeds preferentially on corn, millet, cotton, and sorghum, whereas the *RS* on rice and several pasture grasses (Pashley 1986; Cano-Calle et al. 2015). *CS* and *RS* also have different rates of development and fitness depending on the host plant used (Pashley et al. 1995; Meagher et al. 2004; Busato et al. 2005). They also differ in mating behavior, such as mating allochronism (Meagher et al. 2004; Schöfl et al. 2011; Juárez et al. 2014) and pheromone composition (Groot et al. 2008; Lima and McNeil 2009). A broad study of different populations from Brazil recently recognized several molecular markers and loci under selection when comparing the different strains feeding on different host plants (Silva-Brandão et al. 2018).

Considering the participation of the gut microbiota in the processes of host adaptation to new food resources and of speciation, we hypothesized the gut microbiota may be involved in the process of host-strain adaptation in *S. frugiperda*. To our hypothesis hold true, we predict that the gut microbiota of the strains differs from each other and therefore we also expect a different functional contribution from each other in exploiting similar host plants. In addition, we predicted that alterations in the gut microbiota within one strain from one host plant to another would be less conspicuous than the changes in the microbiota between strains in the same host plant.

Our investigation addressed field-collected insects, which carry a much higher variation in the gut microbiota than those maintained under controlled laboratory conditions (Gomes et al. 2020). Assessing the variation available under field conditions can provide essential information on potential symbionts that could be ecologically important to their hosts in their natural habitats. The selection pressure in natural and laboratory conditions are quite different and can lead to the selection of distinct traits, including the interaction with symbiotic bacteria (Paniagua et al. 2018).

## Material and methods

### Sampling

Larvae of the fall armyworm were collected from corn, cotton and millet crops cultivated in the same landscape in the western region of Bahia state, in the district of Roda Velha, Brazil (12°42’0” S 45°50’0” W). In this area there is crop rotation and some of them also cooccur, moreover there is no evidence indicating occurrences of migration of *S. frugiperda* in the area, therefore we assume that the caterpillars collected on the different host plants analyzed in this study correspond to the same population of *S. frugiperda*. The larvae were collected and directly fixed in RNAlater™ (Thermo Fisher), taken to the laboratory where the width of their head capsules was individually measured under a stereoscopic. Only larvae with head capsules in the range of 1.6 – 2.9 mm were used in the experiments.

### *Spodoptera frugiperda* host strains identification

Strain identification was performed using the DNA extracted in the same way described below (session 2.3) from part of the larvae tegument. Then, RFLP-PCR analysis of a partial sequence of the mitochondrial COI gene using the primers set JM76 (5’
s-GAGCTGAATTAGGRACTCCAGG-3′) and JM77 (5′ - ATCACCTCCWCCTGCAGGATC-3′) (Levy et al. 2002).The PCR mixture contained 100-150 ng of DNA, 1.5 mM of MgCl_2_, 1x PCR buffer, 0.2 mM of each dNTP, 0.32 μM of each primer and 0.63U of GoTaq® DNA Polymerase (Promega) in a total volume of 25 μL. The thermocycling condition was 94°C x 1 min (1x) followed by 33 cycles at 92°C x 45 s, 56°C x 45 s, 72°C x 1 min, and a final extension at 72°C x 3 min (1x).

Amplicons were then subjected to restriction analysis using the *Msp*I (HpaII) restriction endonuclease. This enzyme produces two fragments of 497pb and 72pb for amplicons of the *CS*, and no digestion of the 569-bp amplicon for *RS* is observed. We used 10 μL of the PCR product added to 18 μL nuclease-free water, 2 μL 10x Buffer Tango and 1 unit of *Msp*I. Samples were gently mixed, spun down for a few seconds on a tabletop centrifuge and incubated overnight at 37°C. Subsequently, the resulting products of digestion were verified using 1.8% agarose gel electrophoresis following standard procedures (Sambrook and Russell 2001).

### Gut isolation and genomic DNA extraction

Midgut dissection was performed under aseptic conditions under a laminar flow hood after larval surface sterilization in cooled 70% ethanol solution added with 0.2% sodium hypochlorite (5 min). Larvae were washed once in sterile water and transferred to sterile saline solution (125 mM NaCl, 4°C) for Midgut dissection. Tissues were stored in absolute ethanol at -20°C until DNA extraction. The midguts of individual larvae were grouped according to strain and host plant. Three true biological replicates were established for each treatment (1 replicate = pooled guts of 3 larvae). The homogenization was performed in liquid nitrogen.

DNA was extracted using the protocol for genomic DNA preparation from RNAlater™ (Thermo Fisher) preserved tissues with some modifications. The macerate of the pooled midguts was placed in 750 μL digestion buffer (60 mM Tris pH 8.0, 100 mM EDTA, 0.5% SDS). Proteinase K was added to a final concentration of 500 μg/mL and mixed well by inversion. Samples were incubated overnight at 55°C. Afterwards, 750 μL of phenol: chloroform (1:1) was added and rapidly inverted for 2 min. Samples were centrifuged at a toptable centrifuge for 10 min. The aqueous layer was collected and the process of phenol: chloroform extraction was repeated twice before a final extraction with chloroform. The aqueous layer was collected and added to 0.1 volume of 3 M sodium acetate, pH 5.2, and 1 volume 95% ethanol. Samples were mixed by inversion, incubated for 40 min at -80°C, and centrifuged (27,238 *g* x 30 min x 4°C). The pellet obtained was washed twice in 1 mL of 85% ice-cold ethanol, centrifuged for 10 min after each wash and dried at 60°C during 5-10 min in a SpeedVac. Finally, the pellet was resuspended in DEPC water. DNA concentration and quality were estimated using a Nanodrop UV spectrophotometer.

### 16S rDNA sequencing and analysis

DNA samples were PCR-amplified using primers targeting the V3-V4 region of the 16S rRNA gene (Klindworth et al. 2013). The reaction was programmed at 95°C for 3 min (1 cycle), followed by 25 cycles at 95°C for 30 s, 55°C for 30s and 72°C for 30 s, with a final extension (1 cycle) at 72°C for 5 min. DNA samples was sent to the Animal Biology Laboratory (ESALQ / USP, Piracicaba, SP) for Illumina library preparation and sequencing. Genomic DNA was amplified by PCR with primers targeting the hypervariable V3–V4 region of the 16S rRNA gene. Paired-end reads (2x – 250 bp) were generated with multiple barcodes and sequenced on the Illumina MiSeq platform.

The sequences were processed using QIIME2 (2017.4 release) (Caporaso et al. 2010) according to the developers’ recommendations. The demultiplexed reads were quality filtered, had singletons removed, denoised of truncated reads (250 bp length), and filtered for phiX and chimera sequences with the *q2-dada2* plugin. A feature table was generated with a summary of amplicons sequence variants (ASV), a method shown to have higher resolution than the OTU method, with biological meaning independently from a reference database (Callahan et al. 2017). All ASVs were aligned with the *mafft* program (via *q2-alignment* plugin), and the construction of the phylogenetic tree was performed with FastTree. The sequences were rarefied to a depth equivalent to the smaller total count in our samples to maintain all of them in our analysis. Rarefied feature tables were used as input for alpha and beta diversity analyses and statistics using the core-metrics-phylogenetic method. We examined alpha diversity using Shannon and Simpson’s index, both estimator of species richness and evenness, being the first more weight on species richness and the second on species evenness. And Chao1 index, an abundance-based estimator that gives more weight to the low abundance species. For the beta-diversity we used Jaccard distance that does not take the ASVs abundances into account, just the presence and absence in the samples, and Jackknifed weighted UniFrac distance, that considers the phylogenetic relationships and the abundance of the ASVs. We used them as a basis for hierarchical clustering with Unweighted Pair Group Method with Arithmetic Mean (UPGMA) and PCoA plots. The variation degree among the repetitions was evaluated using the Jackknife technique and displayed by confidence ellipsoids around the samples. Pairwise PERMANOVA (999 permutations) was used to detect differences in composition (β-diversity) between groups. Taxonomy was assigned to ASVs using pre-trained (V3-V4) classifier via *q2-feature-classifier* trained on the Silva_132_99_16S.fna file and clustered at 97% similarities (341-806 region, seven-level taxonomy). Functional profiles from 16S rRNA data were predicted using the *Phylogenetic Investigation of Communities by Reconstruction of Unobserved States 2* (PICRUSt2) plugin for QIIME2. For each sample, the composition of The Kyoto KEGG pathway abundances from the predicted KEGG ORTHOLOGY (KO) abundances were made following the default protocols of the PICRUSt2 GitHub page (https://github.com/picrust/picrust2/wiki).

We used the Statistical Analysis of Metagenomic Profiles (STAMP) software (Parks et al. 2014) to identify taxa that differ significantly between groups. White’s non-parametric t-test was used for taxonomic comparisons and the bootstrap method was used to calculate the confidence intervals. Functional comparisons were done using the Welch’s t-test to test statistical hypothesis, Bonferroni for *p*-values correction and Welch’s inverted as a confidence interval method.

## Results

Our sampling effort was adequate to access the diversity of the microbiota in the gut of both strains of *S. frugiperda* as the rarefaction curves based on the Shannon index did not change after sampling 8,000 sequences, much below the near 30,000 sequences obtained for the lowest represented 16S rRNA library (Fig S1).

Alpha diversity was similar between strains when disregarding the host plant in all calculated indices (Fig. 1). When the alpha diversity was compared within the groups of host plants, the corn strain showed higher Shannon and Simpson indices when the caterpillars fed on millet. On the other host plants, the gut microbiota showed similar values of Shannon, chao1 and Simpson indices (Fig. 1). The beta diversity of the gut bacterial communities of the *RS* and *CS* strains were strongly influenced by the host plants tested – corn, millet and cotton, but no differences in the diversity of the gut bacteria between *RS* and *CS* were detected when feeding on the same host plant (Table 1).

**Figure 1.**
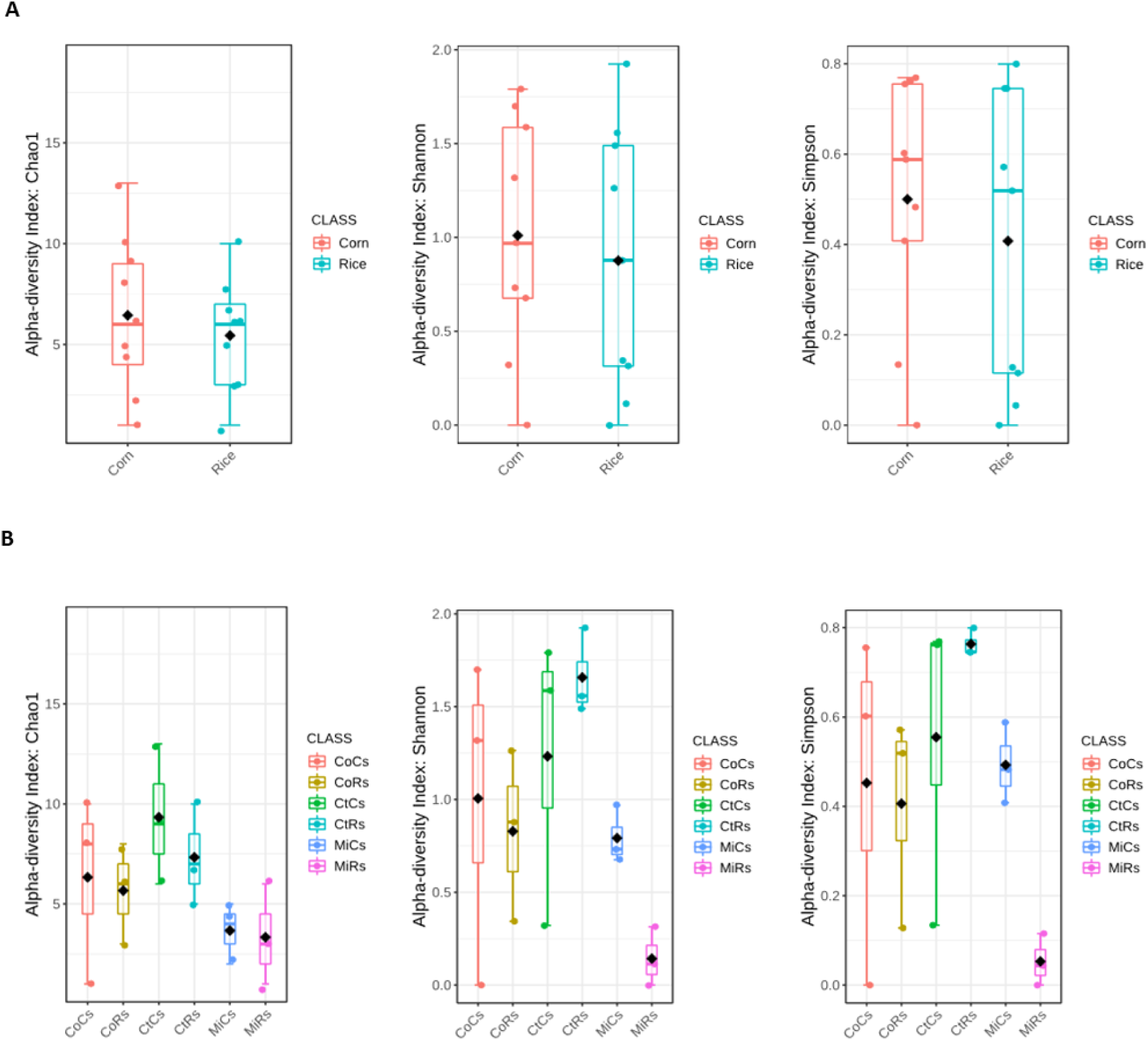
Boxplot comparing the Shannon, Chao1 and Simpson diversity index in samples from *Spodoptera frugiperda* midgut of the corn (CS) and rice strains (RS) regardless diet (A), and considering the diet: corn (Co), millet (Mi) and cotton (Ct).

**Table 1.** Pairwise PERMANOVA comparisons based on weighted UniFrac distance matrices among the different treatments after 999 permutations.

| **Group 1** | **Group 2** | **Sample size** | **pseudo-F** | **p-value** | **q-value** |
| --- | --- | --- | --- | --- | --- |
| **CS - Corn Plant** | **RS – Corn Plant** | 6 | 0.615 | 0.886 | 0.886 |
| **CS – Cotton Plant** | **RS - Cotton Plant** | 6 | 1.126 | 0.284 | 0.525 |
| **CS – Millet Plant** | **RS – Millet Plant** | 6 | 0.329 | 0.816 | 0.874 |
| **Corn** | **Cotton** | 12 | 4.087 | 0.012 | 0.024 |
| **Corn** | **Millet** | 12 | 5.098 | 0.052 | 0.052 |
| **Cotton** | **Millet** | 12 | 9.117 | 0.016 | 0.024 |

Abbreviations: CS= Corn Strain; RS= Rice strain

UPGMA clustering analysis based on the V3-V4 region of the 16S rRNA gene yielded three clusters, each of them grouping most of the samples of each host plant. The replicates of the gut microbiota of the cotton-fed *RS* larvae were the most distinct, all of them resolved in one of the clusters formed. The cotton-fed *CS* replicates grouped with the millet samples and the other with the corn-fed larvae. All corn-fed larvae replicates resolved together except for one *RS* replicate that clustered with the millet replicates. Likewise, the millet replicates were in the same cluster except for one *RS* that clustered with the corn replicates (Fig. 2).

**Figure 2.**
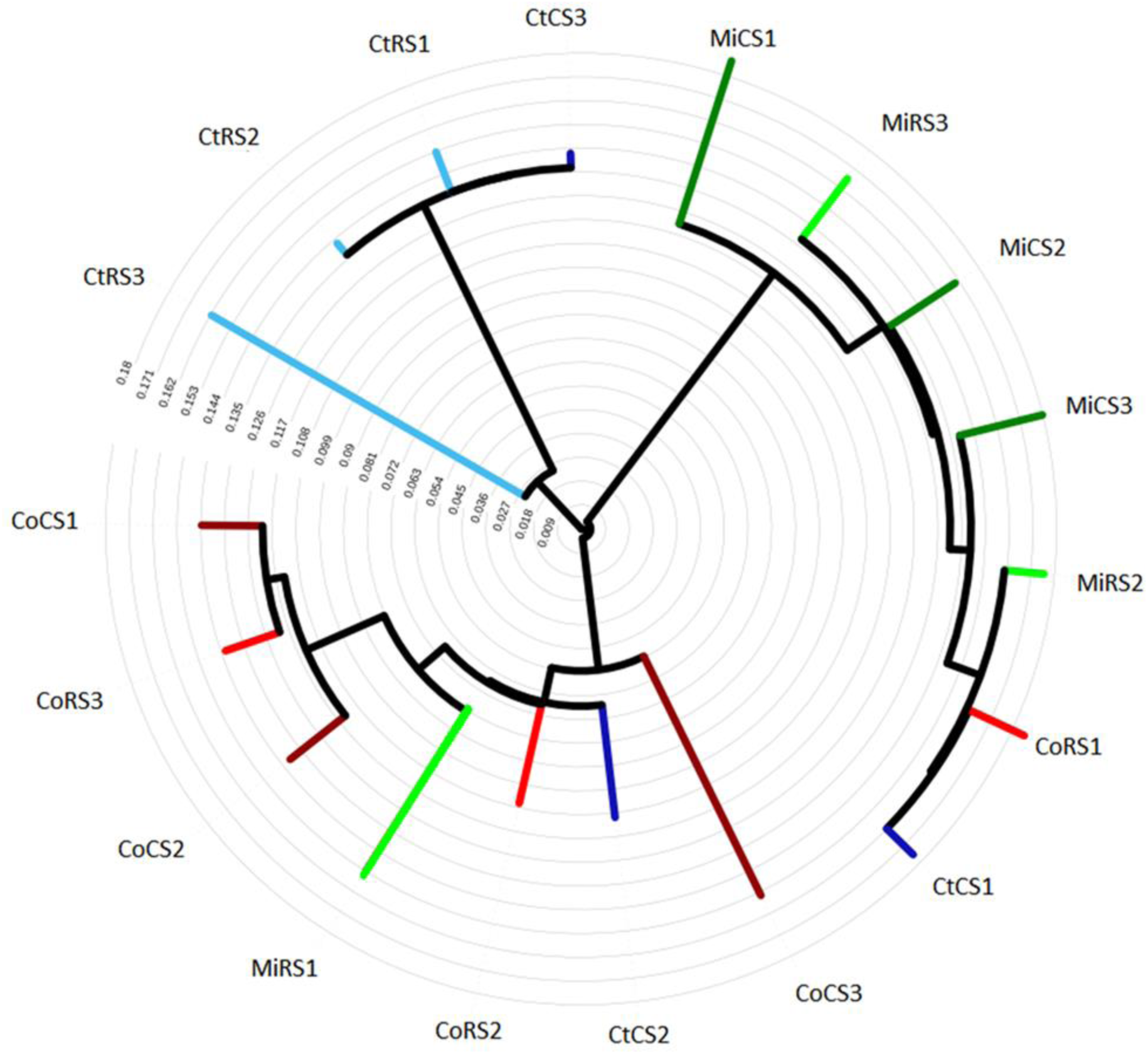
Jackknifed Weighted Pair Group Method with Arithmetic mean (UPGMA) cluster tree of *S. frugiperda* corn strain (*CS*) and rice strain (*RS*) gut microbiota when feeding on corn (Co), millet (Mi) and cotton (Ct).

We identified 17 phyla of bacteria in the midgut of *S. frugiperda* (Fig. S2). The gut microbiota of larvae feeding on the monocots corn and millet was dominated by *Firmicutes*, followed by *Proteobacteria* regardless of the host plant strain (*RS* or *CS*) (Fig. 3). But a significant increase in the relative proportion of *Proteobacteria* was observed in both *RS* and *CS* larvae when feeding on the dicot cotton (Fig. 3). Differences in the gut microbiota composition at the phylum level in between *RS* and *CS* when using the same host plants were observed only in larvae fed on cotton (Fig. 3).

**Figure 3.**
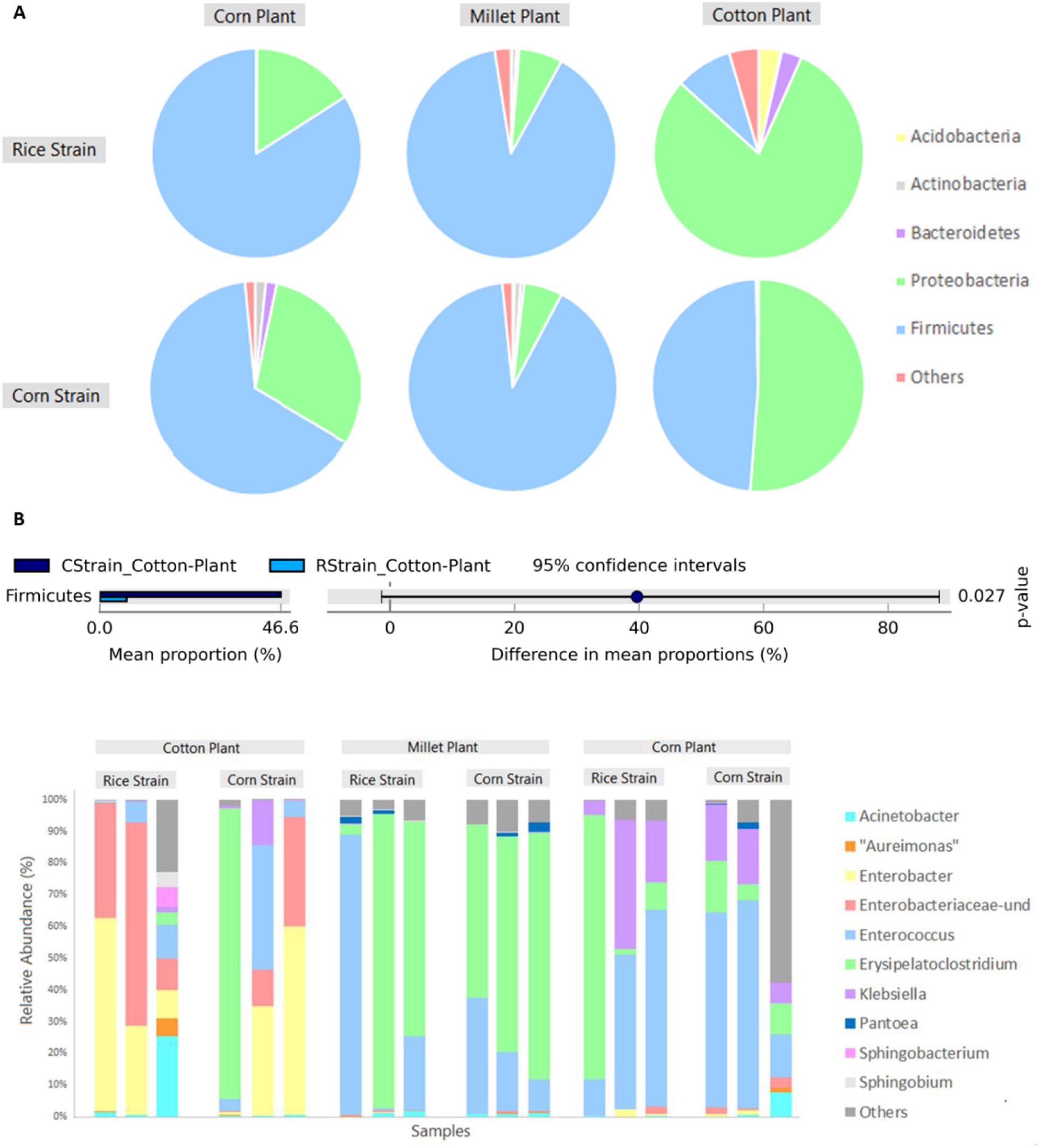
Relative proportion (%) of bacterial phyla abundance between the gut microbiota associated with corn and rice strains of *S. frugiperda* when feeding on corn, millet, and cotton (A) and cotton (B).

*Enterobacter, Enterobacteriaceae, Klebsiella* and *Acinetobacter* (γ-*Proteobacteria, Enterobacteriaceae*) represented most of the diversity of enterobacteria in the midgut of *S. frugiperda. Firmicutes* was mostly represented by *Enterococcus* (*Bacilli, Enterococcaceae*) and *Erysipelatoclostridium* (*Erysipelotrichia, Erysipelotrichaceae*). The unidentified *Enterobacteriaceae* and *Enterobacter* represented most of the diversity of the larval midgut microbiota from cotton; *Enterococcus* and *Erysipelatoclostridium* from millet; and *Enterococcus* and *Klebsiella* from corn (Fig. 4).

**Figure 4.**
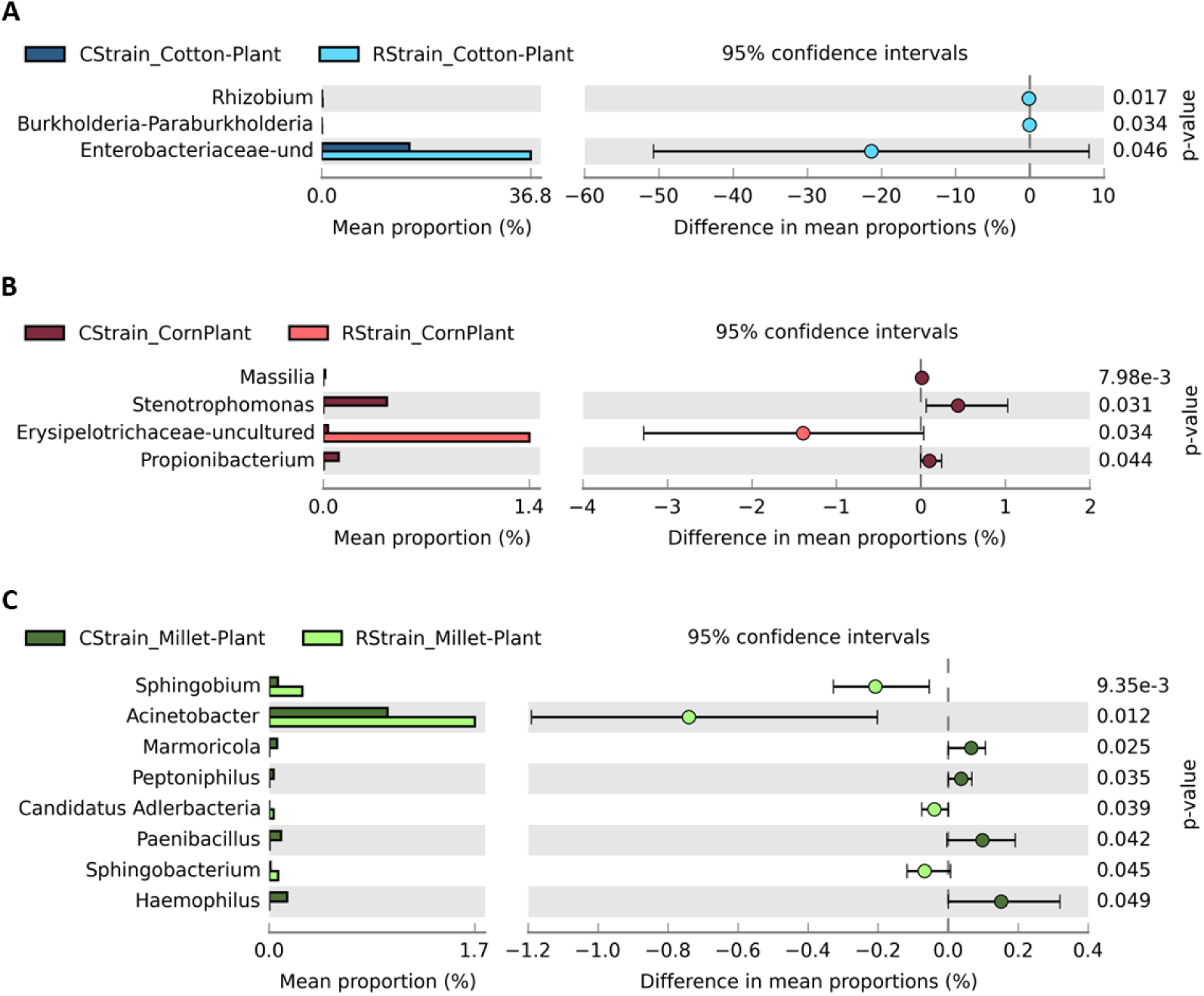
Bacterial genera of the gut microbiota associated with corn and rice strains of *S. frugiperda* when feeding on corn, millet, and cotton.

Comparisons of the relative abundance of the most common ASVs in the midgut of *RS* and *CS* larvae of *S. frugiperda* at the genus level indicated that the *Enterobacteriaceae* was more abundant in *RS* than in *CS*, but only when larvae were collected in cotton (*p* = 0.046). Larvae fed on millet and corn presented differences only in the minor representatives of the gut microbiota. In the corn plant *RS* larvae exhibited a lower relative abundance in *Stenotrophomonas* (*p* = 0.031) and a higher abundance of *Erysipelotrichaceae* (*p* = 0.034) than the *CS* larvae. In the millet plant the abundance of *Sphingobium* (*p* < 0.005) and *Acinetobacter* (*p* = 0.012) in the midgut of *RS* differed from that in the midgut of *CS* larvae (Fig. 5).

**Figure 5.**
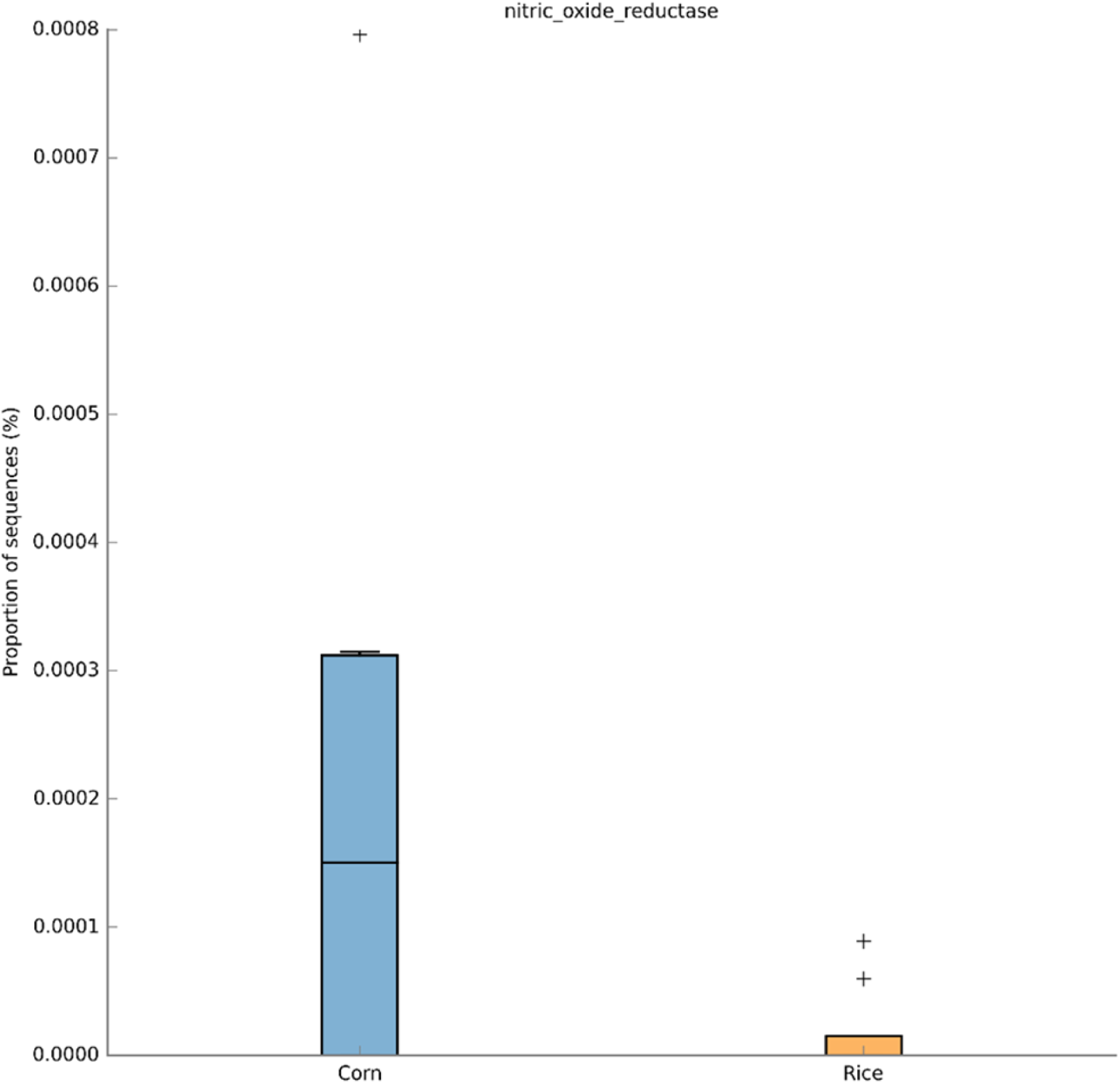
Differences in relative proportion (%) of bacterial genus abundance in the gut microbiota associated with corn and rice strains of *S. frugiperda* when feeding on (A) corn, (B) millet, and (C) cotton.

The small differences in the diversity of the midgut microbiota between the strains were enough to allow the detection of changes in the potential functional contribution of the gut microbiota of *RS* and *CS*. But although highly significant, the variation in the proportional changes of the potential functional contribution was very minor, and possibly with no physiological impact. Differences in energy metabolism between *CS* and *RS* were detected but only for potential contribution of the *norF* gene, which belongs to a gene cluster involved in nitrogen metabolism and microbial defense against nitric oxide toxicity (p=0.045). The gut microbiota of the *CS* had a higher potential contribution for the *norF* protein (Fig. 6). Pairwise comparisons of *CS* and *RS* in each host plant detected that the midgut microbiota of the *RS* had a higher potential contribution to metabolism (*wcaE* - glycosyltransferases, beta-glucuronidase - secondary metabolites) and the processing of genetic information (integrase) in millet, while the contribution of the gut microbiota of the *CS* was much higher to the signaling and cellular processes (*hpaX, fliJ, hoxN, flgL, dmsB, fliS, fliH, cheV, flgE, fliM, flgD, fliNY, fliE, fliI, fliP*) (Fig. 7). The potential contribution of the gut microbiota to signaling and cellular processes (*pilX, terZ, TC*.*AAA*) was also higher in the *CS* than the *RS* strain when feeding on corn. In cotton, the gut microbiota of both host-adapted strains had increased contribution to metabolism, but while the contribution of the microbiota of the *CS* to metabolism was due to the higher abundance of glyceride 2-quinase (carbohydrate metabolism) and L-serine dehydrate (amino acid metabolism), the potential contribution of the gut microbiota of the *RS* to metabolism was due to the increased abundance of oxidases (glycine and vitamin cofactors) and amylosucrase (carbohydrate metabolism). The gut microbiota of the *RS* also had a higher potential of contribution to the processing of genetic information (*pinR*) and signaling and cellular processes (ATP-binding cassette) than that of the *CS* strain when feeding on cotton (Fig. 7).

**Figure 6.**
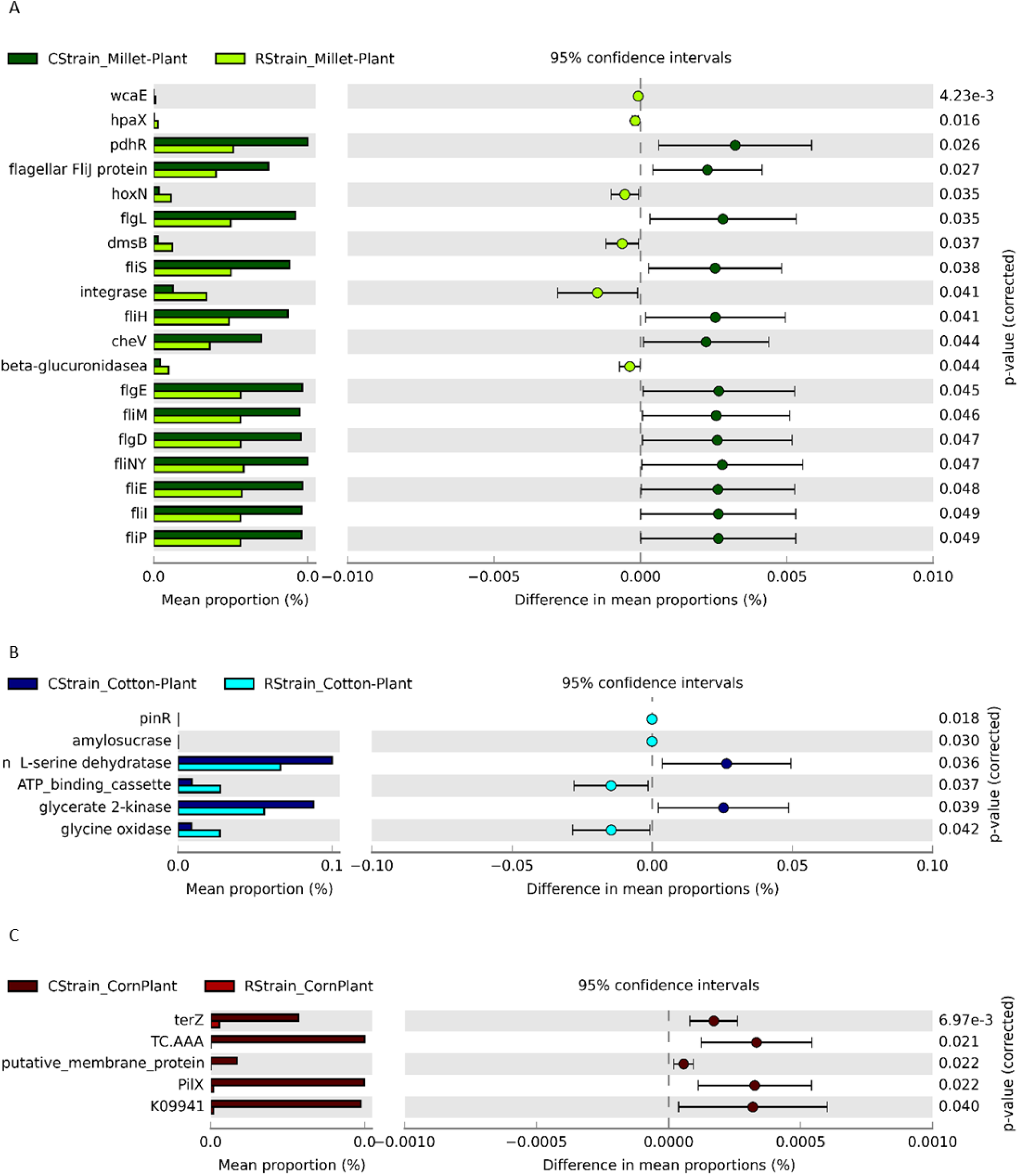
Differences in PICRUSt functional prediction between the gut microbiota associated with corn and rice strains of *S. frugiperda* regardless of host plant.

**Figure 7.**
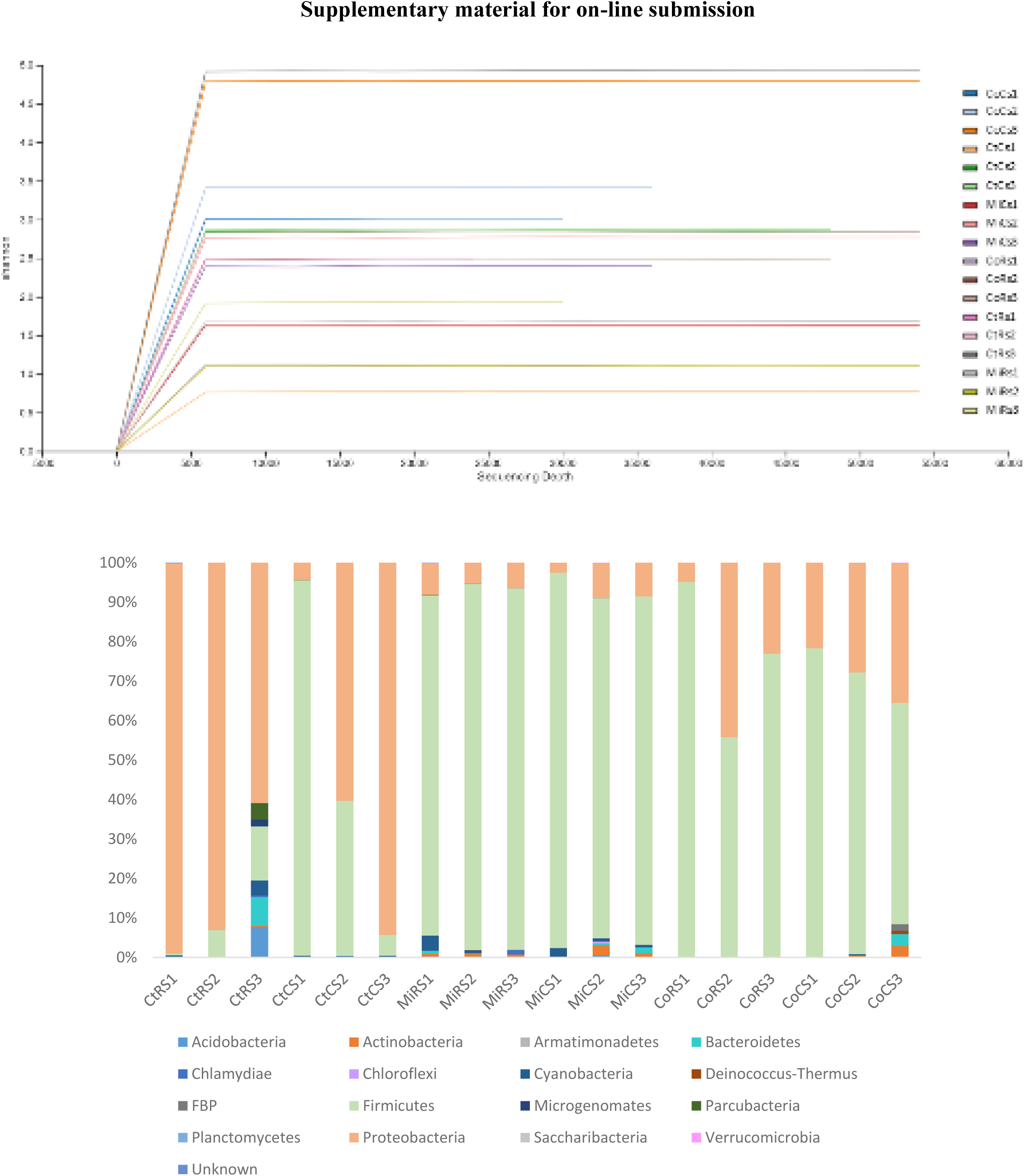
Difference in relative proportion (%) of PICRUSt functional prediction of the gut microbiota associated with corn and rice strains of *S. frugiperda* when feeding on (A) millet (B), and cotton, (C) corn.

## Discussion

The *CS* and *RS* strains of *Spodoptera frugiperda* share an overall similar microbial diversity when feeding on the same host plants, although there is a significant genetic divergence between the host-adapted strains. The midgut microbiota of *CS* and *RS* larvae went through similar changes when the larvae were feeding on the different host plants analyzed. These results suggest that the diet, but not the strains play a more important role in shaping the gut bacteria community structure of these larvae. These findings are like those found in humans, where host genetics played a minor role in determining microbial composition than environmental factors such as diet (Rothschild et al. 2018).

There are convincing molecular and biological data demonstrating the *RS* and *CS* strains of *S. frugiperda* are in a process of speciation and host plant adaptation (Dumas et al. 2015; Gouin et al. 2017). Although we can clearly detect a shift in the midgut microbiota in response to the different host plants tested, it was not possible to detect selection of specific bacteria species that would characterize the diversity of the gut microbiota of each strain.

Thus, we first argue that the lack of a clear difference in the midgut microbiota of *RS* and *CS* could be explained by the recency of their speciation process (Kergoat et al. 2012). In fact, the strains are still interbreeding and producing viable offspring (Dumas et al. 2015), although postzygotic mechanisms of reproductive isolation are acting as facilitators of speciation in *S. frugiperda* (Kost et al 2016). Therefore, the short evolutionary period may not be enough for selection of specific members of the gut microbial community by the host. In addition, the gut microbiota composition is regulated by several factors, such as immune system (e.g., lysozyme, reactive oxygen species and antimicrobial peptides) (Chapelle et al. 2009), physical and chemical properties of the gut (e.g., pH and redox conditions), presence of digestive enzymes and development of specialized structures in the gut. Perhaps these processes that are involved in controlling and selecting the components of the gut microbiota have not differentiated enough between the strains to cause differences in the midgut bacterial community of both strains.

Alternatively, we also argue that the existing differences among bacterial members of the midgut of each strain could not be adequately accessed using the 16S metabarcoding procedure based on short sequences of the 16S rRNA gene. Gut symbionts become host specialized during the process of host evolution (Frese et al. 2011; kwong et al. 2014) and data from single cell genome sequencing have demonstrated the existence of significant genomic differences even within bacterial cells that share highly homologous 16S rDNA sequences (Engel et al. 2014).

Our third argument is based on the presence of individuals from mixed crossings from *CS* and *RS*, as our field samples also had low *RS* larvae representation (corn=28%; millet: 25%; cotton:15%; data not shown). As *RS* x *CS* crossings naturally occur in the field (Nagoshi and Meagher 2003), and the bacterial composition of the gut of hybrids can differ from that of parentals (Miller and Miller 1996), we cannot rule out that the existence of *CS-RS* or *RS-CS* hybrids in our samples may have diluted the differences between *RS-RS* and *CS-CS* midgut bacterial communities.

Finally, we examined only the larval midgut in view of its important function in food digestion and assimilation (Dow 1987; Billingsley 1996). However, this gut region is predisposed to be an adverse environment for microorganisms due to enzymes, antimicrobial peptides, and high alkaline pH (Dow 1987). In addition, another possible obstacle bacteria face to colonize the midgut of lepidopterans is the presence of a type 1 peritrophic membrane, which is produced and continuously replaced, moving subsequently with the food bolus along the digestive tract (Wu et al. 2016; Hegedous et al. 2019). Perhaps some resident symbionts could colonize the hindgut, which is known to be more favorable to bacteria due to the lack of digestive enzymes, presence of favorable morphological structures, and of ions and metabolites released in the urine (Douglas 2015).

The midgut of *CS* and *RS* larvae of *S. frugiperda* carries basically the same group of bacteria in each host plant analyzed. *Erysipelatoclostridium, Enterococcus*, one unidentified genus of *Enterobacteriaceae* (*Enterobacteriaceae*-und), *Klebsiella* and *Acinetobacter*. The relative abundance of these bacteria was altered depending on the host plant, but changes observed in each host plant were quite similar between *RS* and *CS* larvae.

*Enterobacteriaceae*-und and *Enterobacter* were the most abundant in the gut microbiota of cotton-fed larvae. *Enterobacter* is commonly found in the gut and was shown to reproduce in the insect midgut (Watanabe et al. 2000; Tanada and Kaya 2012).

*Erysipelatoclostridium*, the most abundant in the bacterial community of the midgut of larvae from millet is reported as one of the major genera in the human gut, where it utilizes proteins and saccharides as substrates, producing acetate, carbon dioxide, hydrogen, formate and lactate (Oliphant and Allen-Vercoe 2019). Food-induced changes in the *Firmicutes*: *Bacteroidetes* ratio in the gut of humans were noticed by alterations in the abundance of *Erysipelatoclostridium* (Smith-Brown et al. 2016). In model vertebrate animals, *Erysipelatoclostridium* has been associated with the upregulation of glucose and fat transporters in the gut (Günther et al. 2007). Increased abundance of *Erysipelatoclostridium* in the gut microbiota of rats has been positively correlated with the fatty acid isovalerate (by-product of leucine fermentation), affecting the digestibility of the food source (Han et al. 2018). In insects, *Erysipelatoclostridium* has been reported as an unculturable gut symbiont of field populations of *Spodoptera litura*, but nothing is known on the role of this bacterium in the gut of insects (Yalashetti et al. 2017).

Larvae of *S. frugiperda* fed on corn present a high abundance of the Gram-negative bacterium *Klebsiella* sp. This genus has been isolated from corn leaves and has also been identified in the gut of many lepidopteran species, demonstrating the ability of *Klebsiella* to colonize their digestive tract (Chen et al. 2016; Snyman et al. 2016). In addition, *Klebsiella* can play a role as a mediator of insect-plant interactions. The presence of this bacteria in oral secretions of *S. frugiperda* larvae can regulate the expression of the herbivorous induced proteinase inhibitor gene (*mpi*) in corn (Acevedo et al. 2017).

*Enterococcus* was present in all our samples but with higher abundance in samples from corn and millet. Nearly 40 species of *Enterococcus* are known and are predominantly reported as commensals of the gastrointestinal tract (Ramsey et al. 2014). Studies with *S. littoralis* suggested the existence of a clear symbiotic relationship with *Enterococcus mundtii*, a biofilm-like structure maker that contributes with the secretion of antimicrobial peptides (AMP) supposedly contributing to the host as an additional chemical barrier against pathogens (Shao et al. 2014, 2017).

Another aspect that we observed in our study was the importance of diet in shaping the midgut microbiota as already documented in the literature (Gayatri Priya et al. 2012; Tang et al. 2012; Yun et al. 2014). Our samples were collected from a single, well-defined landscape; thus, we did not expect much variation in the genome of the sampled insect population, a factor that could also interfere with the gut microbiota. Therefore, variation lies mainly in the host plant and strain. Analyzing the gut microbiota composition, we observed a dysbiosis in the gut driven by diet. These changes in the community structure may benefit *S. frugiperda* as an agriculture pest. One example of this is the case of the western corn rootworm, in which the gut microbiota shifted in response to crop rotation, increasing the abundance of *Klebsiella* and *Stenotrophomonas*. This change in the microbiota led to an increase of bacterial enzymes in the gut that aided in food digestion, as these microbial enzymes were insensitive to soybean cysteine protease inhibitors (Chu and Mazmanian 2013).

The analysis of the potential functional contribution of the midgut microbiota of *S. frugiperda* resulted in only a single difference between the strains when we disregard the host plant. The higher potential contribution of the *CS* midgut bacteria to denitrification due the higher abundance of species carrying putative *norF* genes, which has been described as part of a cluster of genes encoding nitric-oxide reductases. Analysis of mutants of this cluster of genes indicated *norF* is involved in the regulation of nitric oxide reductase activity (de Boer et a. 1996). Differences in the potential functional contribution between bacterial communities arise either by the presence of different bacteria members or the exposure to diverse environmental conditions. Since we did not find differences in the composition of the bacterial communities of the midgut between strains when we disregarded the host plant, we can assume that changes detected in the potential functional contribution of the midgut microbiota is likely a result of different biochemical conditions in the gut of each strain. Environments with low oxygen concentration allows the expression of genes involved in the process of denitrification, enhancing the bacterial survival and growth capability in anaerobic environments (Delgado et al. 2007).

The major facilitator transporter 4-hydroxyphenylacetate permease (*hpaX*) is a transmembrane transporter of 4-hydroxyphenylacetate (HPC), which is the first product of the degradation process of 4-hydroxyphenyl-acetic acid (HPA). HPA has been linked to the overgrowth of bacteria in the gut (Chalmers et al. 1979). HPC is also known to be a fermentation product of amino acids and could be anaerobically degraded by denitrifying bacteria (Seyfried et al. 1991). The dimethyl sulfoxide reductase subunit B (*dmsB*) is a subunit of the terminal anaerobic electron transfer enzyme DMSO reductase, which is also involved in anaerobic metabolism. The high affinity nickel permease (*hoxN*) is involved in the incorporation of nickel into hydrogenase and urease enzymes. The acquisition, delivery, and incorporation of nickel into target enzymes (e.g., urease and hydrogenase) are essential for the catalytic activity of nickel-dependent enzymes, and some bacteria, such as *Escherichia coli, Helicobacter pylori, Yersinia* species, *Salmonella, Shigella* and *Mycobacterium tuberculosis* rely on the system of nickel trafficking for their survival and pathogenicity (Degen and Eitinger 2002; Mulrooney and Hausinger 2003; Higgins et al. 2012).

The midgut microbiota of *CS* fed on millet had a higher number of genes involved in motility and host colonization such as *fli* and *flg*, which encodes for flagellar proteins, as well as *cheV*. Some of these genes (*flgD, flgE, flgL, fliI, fliM, fliP*) are common to most bacterial taxa, while others (*fliE, fliJ, fliS, fliH*) are sporadically distributed (Liu and Ochman 2003). Besides the expected contribution of bacterial flagellum to movement, flagellum also affects cell adhesion, biofilm formation and host invasion (Macnab 2003). *cheV* is a chemotactic protein consisting of a *cheW* domain fused to a phosphorylatable receiver domain. Evolutionary genomics studies suggested *CheV* as an additional adaptor for the accommodation of specific chemoreceptors within the chemotaxis signaling complex (Ortega and Zhulin 2016). Bacterial chemotaxis is fundamental to allow bacteria to detect and follow chemical gradients in their environment (Baker et al. 2006). Therefore, taking all this together, there is a possibility that the *CS* has a microbiota with greater capacity to colonize the host midgut than the *RS*.

In samples from cotton, the *CS* bacterial community contain species commonly encoding higher levels of L-serine dehydratase, that was shown to be essential for colonization of the avian gut by *Campylobacter jejuni* (Velayudhan et al. 2004). Additionally, the gut microbiota of *CS* contributes with higher levels of glycerate-2-kinase that degrade glucose via a nonphosphorylative Entner-Doudoroff pathway, a pathway central for energy and carbon metabolism (Conway 1992).

Finally, the genes predicted for the bacterial community in the midgut of larvae feeding on corn were predominantly related to signaling and cellular processes (*TC*.*AAA, tellurium resistance protein-TerZ, pilX*). For all these predicted genes, *CS* had a higher abundance, suggesting a microbial community better suited to communicate with its environment and/or to better respond to temporal variations of external signals they experience.

We understand our study may have certain limitations, such as the low number of biological replicates. But this limitation was compensated by using pooled samples from the field and by adopting very stringent statistical analysis and care in data interpretation, particularly in cases with outliers. We also understand that the functional analysis provided brings certain limitations since it is based on the detection of 16S rRNA gene amplicons, and bacteria detection by DNA amplification cannot be considered as direct evidence that the taxa amplified are metabolically active. Finally, although Picrust is useful for predicting the potential functional contributions of the gut microbiota, the use of this tool brings limitations that should be highlighted such as those associated with the availability of appropriate references, biased primers, and gaps or inaccuracies in pathway annotation or gene function assignments. Future work to address these limitations could test the real contribution of the gut microbiota of *S. frugiperda*.

In conclusion, our data demonstrate that the larval midgut of *S. frugiperda* harbor a bacterial community that varies according to the host plant. We also demonstrate that the midgut bacterial community consisted predominantly of *Firmicutes* followed by *Proteobacteria* when the larva feeds on corn and millet, with an opposite pattern when the larva feeds on cotton, regardless of the host strain of *S. frugiperda*. Differences at the genus level between the bacterial community of the *CS* and *RS* and predicted functional groups of low abundance were also detected. Studies of the gut microbes of this important agricultural pest can provide new knowledge not only for their control, but also for a better understanding of processes of host adaptation and evolution in insects.

## Acknowledgements

We are grateful to the São Paulo Research Foundation (FAPESP) (process 2011/50877-0) and the Ministry of Science, Technology, and Innovation (Conselho Nacional de Desenvolvimento Científico e Tecnológico – CNPq: process 403851/2013-0 and 462140-2014/8) for providing funds to this research. We also thank FAPESP for the PhD student fellowship (2017/24377-7) provided to the first author. This manuscript is one of the chapters of the PhD Thesis of the first author.

